# Drug repurposing reveals Posaconazole as a CYP11A1 inhibitor enhancing anti-tumour immunity

**DOI:** 10.1101/2025.02.21.639414

**Authors:** Jhuma Pramanik, Sanu Korumadathil Shaji, Megan Zaman, Bethany Brown, Baojie Zhang, Yumi Yamashita-Kanemaru, Natalie Z. M. Homer, Hosni Hussein, Qiuchen Zhao, Klaus Okkenhaug, Rahul Roychoudhuri, Abhik Mukhopadhyay, Bidesh Mahata

## Abstract

Steroid hormones regulate cell physiology and immune function, with dysregulated steroidogenesis promoting cancer progression by supporting tumour growth and suppressing anti-tumour immunity. Targeting CYP11A1, the first and rate-limiting enzyme in steroid biosynthesis, has shown promise in cancer therapy, but safe and effective inhibitors remain an unmet need. Undertaking *in silico* structure-based drug repurposing approach, we found Posaconazole as an inhibitor of CYP11A1. The docking pose analysis showed that Posaconazole can form multiple hydrogen bonds and hydrophobic interactions with the key residues at the binding site and the cofactor, stabilising the protein-ligand complex. We validated its inhibition efficiency in cell-based assays. In a mouse model of lung metastasis, we demonstrated that Posaconazole restricts metastatic cancer growth by stimulating anti-tumour immunity. These findings highlight Posaconazole’s potential as a research tool to study steroidogenesis and as a candidate for further preclinical and clinical evaluation in pathological conditions associated with local steroid production, such as steroidogenic tumours.

## Introduction

CYP11A1, also known as P450 side chain cleavage enzyme or P450scc, is a crucial member of the cytochrome P450 family. This enzyme catalyses the initial and rate-limiting step in steroidogenesis by converting cholesterol to pregnenolone (Miller and Auchus, 2011; Tuckey, 2005). Pregnenolone serves as the first bioactive steroid in the steroid biosynthesis pathway and acts as a precursor for all other steroid hormones (Schiffer et al., 2019). The conversion of cholesterol to other steroid hormones is termed as steroidogenesis, and sometimes this has been denoted as de novo steroidogenesis to distinguish it from the local conversion of steroids from one species to another (Chakraborty et al., 2021). CYP11A1 is present in the inner mitochondrial membranes of steroidogenic cells (Miller and Auchus, 2011; Tuckey, 2005). Steroid-producing cells are commonly found in the adrenal gland, gonads, and placenta, which are the major contributors to the systemic level of steroids in the body (Holst et al., 2004; Miller and Auchus, 2011; Tuckey, 2005). However, extra-glandular steroidogenesis, also known as local steroidogenesis, is reported in various tissues such as the brain (Belelli and Lambert, 2005; Miller and Auchus, 2011), skin (Slominski et al., 2013), thymus (Vacchio et al., 1994), lung (Hostettler et al., 2012), intestine (Ahmed et al., 2019; Bouguen et al., 2015) and adipose tissues (Li et al., 2015). Interestingly, several types of immune cells have also been identified to produce steroids(Acharya et al., 2020; Chakraborty et al., 2021; Jia et al., 2013; Mahata et al., 2020; Mahata et al., 2014; Wang et al., 2013).

Steroid hormones play crucial roles in diverse physiological processes, including cellular metabolism regulation, electrolyte balance, reproductive organ differentiation, pregnancy maintenance, and immune cell function modulation(Chakraborty et al., 2021). The immunosuppressive properties of steroids have been extensively investigated and utilized clinically for treating various immune-mediated disorders.(Coutinho and Chapman, 2011). Recent studies have demonstrated that the presence of steroids in the tumour microenvironment (TME) promotes cancer progression and metastasis by suppressing anti-tumour immune responses(Acharya et al., 2020; Khadka et al., 2023; Mahata et al., 2020; Taves et al., 2023). Notably, steroid production by immune cells within the TME contributes to the remodelling of the local environment, leading to suppressed immune surveillance and facilitating cancer cell immune evasion(Acharya et al., 2020; Mahata et al., 2020). Understanding the fundamental mechanisms of tumour-induced immunosuppression within the TME is critical for developing novel therapeutic strategies in cancer immunotherapy(Hegde and Chen, 2020). Genetic and pharmacological studies targeting CYP11A1, a key enzyme in steroidogenesis, have suggested the potential of modulating this pathway to reinvigorate anti-tumour immunity^10,12,16^. A substantial body of experimental evidence supports the occurrence of local (extra-adrenal) steroid production within tumours, either through de novo steroidogenesis or steroid conversion. This phenomenon has been observed in various malignancies, including cutaneous, gastrointestinal, prostate, and bone metastatic cancers, among others (Bennett et al., 2012; Bouguen et al., 2015; Ergang et al., 2021; Sandor et al., 2024; Slominski et al., 2013; Taves et al., 2023). Beyond oncology, elevated steroidogenesis has been implicated in other pathological conditions, such as peanut allergy, where CYP11A1 inhibition has been proposed as a potential therapeutic approach(Wang et al., 2013; Wang et al., 2020). However, the development of a safe and efficacious CYP11A1 inhibitor remains an unmet clinical need. The identification of a potent steroidogenic inhibitor could have wide-ranging applications, as small molecule inhibition of CYP11A1 could effectively target the de novo steroid synthesis pathway in various pathological conditions. Given the multifaceted potential of CYP11A1 inhibition, this study aims to identify a novel CYP11A1 inhibitor that can serve both as a valuable research tool for investigating steroidogenesis and as a promising candidate for future therapeutic development.

In this study, we employed a drug repurposing strategy to identify novel CYP11A1 inhibitors, leveraging the efficiency and cost-effectiveness of structure based virtual screening. We utilized the GOLD docking suite to screen three subsets of compounds from the ZINC15 database: FDA-approved drugs, drugs in global use, and compounds under clinical investigation. This approach aimed to overcome the time delays, high attrition rates, and substantial resource commitments associated with traditional drug discovery methods. Our screening process identified Posaconazole, an FDA-approved antifungal drug, as a potent inhibitor of CYP11A1 activity. In vitro studies demonstrated that Posaconazole effectively attenuates pregnenolone synthesis in de novo steroidogenic T helper 2 (Th2) cells, outperforming Aminoglutethimide, the current gold standard inhibitor of steroidogenesis. To evaluate the in vivo efficacy of Posaconazole, we employed a mouse lung metastasis model using B16-F10 metastatic melanoma cells. Our results showed that Posaconazole treatment significantly inhibited tumour progression by reinstating anti-tumour immunity. Specifically, Posaconazole increased the number of anti-tumoral T cells and M1 macrophages while reducing pro-tumoral M2 macrophages in the tumour microenvironment. These findings highlight the potential of repurposing Posaconazole for the therapeutic inhibition of steroidogenesis in pathological conditions associated with elevated steroid production. Moreover, our study provides a valuable tool for biological research, offering a means to inhibit the first and rate-limiting step of the steroid biosynthesis pathway. This advancement will enhance our analytical understanding of de novo steroidogenesis and facilitate further investigations into steroid-dependent processes in various physiological and pathological contexts.

## Results

### Virtual screening and selection of compounds for *in vitro* evaluation

A brief overview of the screening methodology is represented in **Figure 1A**. Four structures of CYP11A1 from human are available in The Protein Data Bank in Europe (PDBe) with IDs. 3N9Y, 3NA0, 3NA1 and 3N9Z, (Strushkevich et al., 2011). The Resolution, R factor and R_free_ values for all these structures were compared (**Supplementary Table 1**). All the structures were compared for clashes or other potential model building errors within the active site residues. 3N9Y structure had clashes at S352, V353 and I461, whereas 3NA1 had a sidechain outlier at S352 and a class with the Q377 residue indicating model building errors near the active site. 3NA0 possessed more fit-to-density (RSRZ) and Ramachandran outliers. 3N9Z was selected and used for all docking experiments. 3N9Z has 22-hydroxycholesterol (22-HC) (PDB chem comp id HC9) as the co-crystallised ligand. Analysis of binding pockets, using SuperStar, of 3N9Z showed both hydrophilic and hydrophobic regions (**Supplementary Figure S1A**,**B**). Aliphatic carbon was used to probe the hydrophobic regions, whereas alcohol carbon was used to find hydrophilic regions.

**Figure 1.**
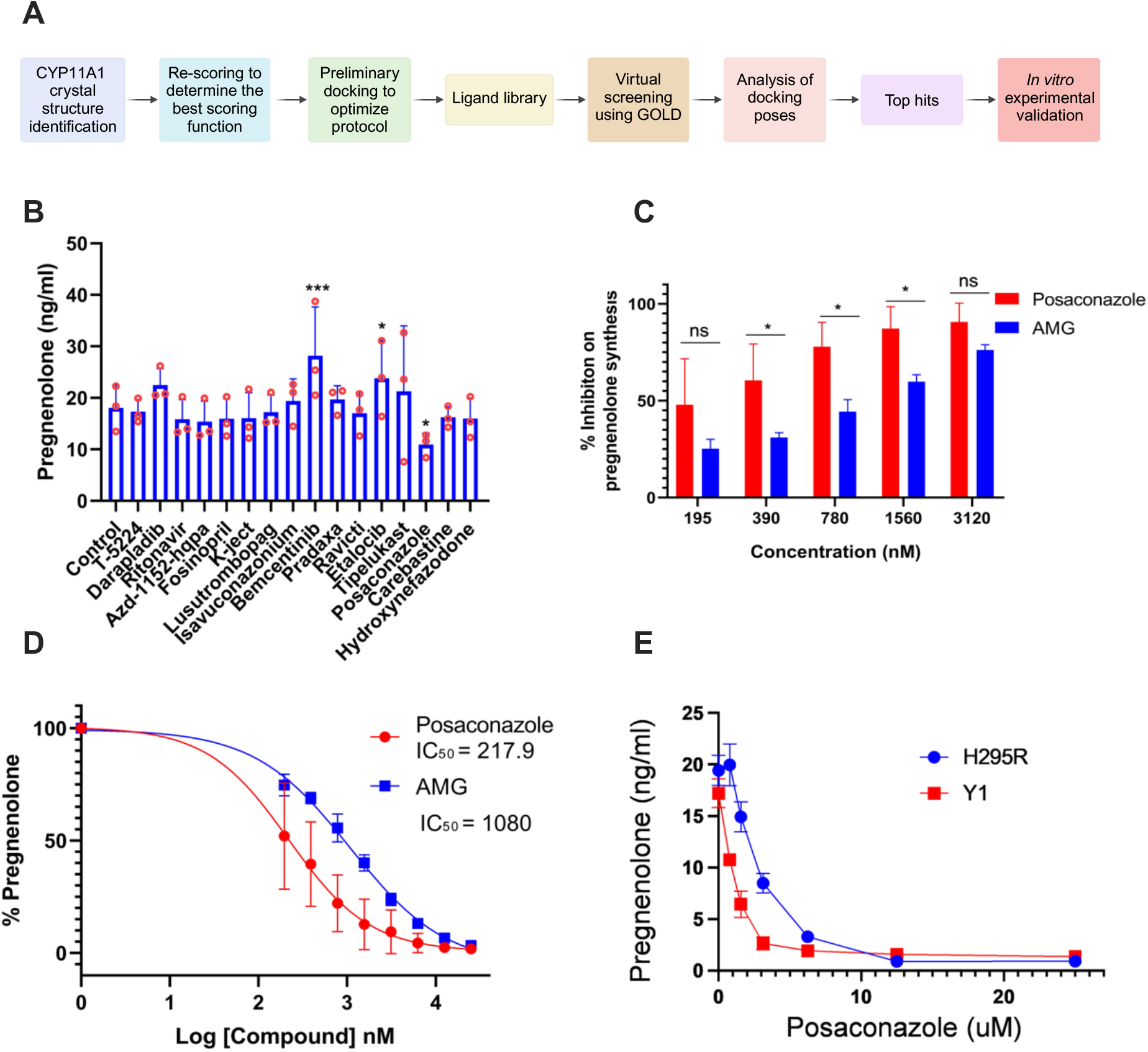
*In vitro* validation of top hits and identification of Posaconazole as a potent inhibitor of CYP11A1. **A**. Schematic of the pipeline used for the identification and validation of CYP11A1 inhibitors using *in silico* drug repurposing approach. **B**. Quantification of Pregnenolone synthesis by Th2 cells after treatment with 1µM of shortlisted drugs. In vitro generated Th2 cells were cultured in the presence of different drugs for 48 hours in charcoal stripped serum containing media. Pregnenolone concentration was measured by ELISA (N= 3, One-Way ANOVA with Fisher’s LSD test, * P ≤ 0.05, ** P ≤ 0.01, *** P ≤ 0.001). **C**. Dose dependent inhibition of pregnenolone synthesis in the presence of Posaconazole and Aminoglutethimide. In vitro generated Th2 cells were treated with different concentration of Posaconazole and Aminoglutethimide for 48 hours and pregnenolone concentration was measured by ELISA (N= 3, Two-Way ANOVA with Sidak multiple comparison, * P ≤ 0.05, ** P ≤ 0.01, *** P ≤ 0.001). **D**. IC50 calculation of CYP11A1 inhibition by Posaconazole and Aminoglutethimide. **E**. H295R and Y1 cells were culture in the presence of increasing concentration of Posaconazole and Pregnenolone concentration in the cell culture supernatant was measured by quantitative ELISA.

After virtual screening of the 7,190 compounds from the above mentioned libraries from ZINC-15, a total of 19 compounds (**Supplementary Figure S1C**) were shortlisted for further experimental analysis. In this final list, we have carefully included compounds that can form polar and non-polar interactions based on the SuperStar analysis of CYP11A1 binding pockets, where both hydrophilic and hydrophobic regions were observed (**Supplementary Figure S1A, B**).

### *In vitro* validation of shortlisted CYP11A1 inhibitors identifies Posaconazole as a potent inhibitor of steroidogenesis

Th2 cells express Cyp11a1, which converts cholesterol to pregnenolone(Mahata et al., 2020; Mahata et al., 2014). However, they do not express downstream enzymes that can catalyse pregnenolone to other steroids (Mahata et al., 2020). Therefore, Th2 lymphocytes are ideal to test Cyp11a1 inhibition by measuring pregnenolone concentration. To shortlist the drugs with any Cyp11a1 inhibitory effect, *ex vivo* differentiated murine Th2 cells were treated with 10 µM drugs for 48 hours, and the inhibitory effect was measured by pregnenolone ELISA (**Supplementary Figure S1D**). Next, to compare the inhibitory efficiency of the selected drugs with Aminoglutethimide, 1 µM of the shortlisted compounds were added to the Th2 culture for 48 h and pregnenolone levels were quantified (**Figure 1B**). The dose was selected based on preliminary titration experiments in this study where Aminoglutethimide demonstrated Th2 steroidogenesis inhibition at sub-micromolar concentrations (Figure 1C). The ELISA analysis identified strong inhibition of Pregnenolone synthesis in Th2 cells treated with Posaconazole (**Figure 1B**). The drugs Oprozomib, Octenidine and Itraconazole showed 94.12 %, 57.55 % and 86.58 % cytotoxicity in Th2 cells, respectively (**Supplementary Figure S1D**). Therefore, these compounds were excluded from further downstream analysis. Posaconazole inhibited the pregnenolone synthesis without any significant reduction in cell viability (**Supplementary Figure S1E**). This result indicated the successful identification of a CYP11A1 inhibitor using the drug repurposing pipeline established to perform the current study. Further *in vitro* and *in silico* analyses were performed on Posaconazole to characterise its potential to inhibit CYP11A1 activity. Since Posaconazole showed a reduction in pregnenolone synthesis without significant change in the cell viability, a dose-dependent analysis of Posaconazole on CYP11A1 inhibition was performed using pregnenolone ELISA. Th2 cells were treated with Posaconazole and Aminoglutethimide with concentrations ranging from 195 nM – 3120 nM for 48 h. The analysis showed a dose-dependent reduction in Pregnenolone levels, indicating an inhibition in CYP11A1 activity (**Figure 1C**). The IC_50_ value for the Posaconazole was found to be 217.9 nM which was four times lower than that of Aminoglutethimide, which showed an IC_50_ of 1080 nM (**Figure 1D**). Aminoglutethimide is the gold-standard CYP11A1 inhibitor and widely used in research. The GOLD score of Aminoglutethimide obtained in the virtual screening was 60, whereas Posaconazole scored 112.4 (**Supplementary Table 2**), which is also in line with the inhibitory activity observed *in vitro*. These observations indicate that Posaconazole is a potent inhibitor of CYP11A1 activity which outperforms the well-established CYP11A1 inhibitor Aminoglutethimide in preventing pregnenolone synthesis in Th2 cells.

### Multiple interactions stabilise Posaconazole in CYP11A1 binding pocket

Posaconazole’s CYP11A1 inhibitory potential observed in the in vitro experiments can be explained by its interactions in the docking pose obtained in the virtual screening (Figure 2). The ligand-protein complex is stabilised by multiple hydrogen bonds and hydrophobic interaction. The bulky nature of Posaconazole fits very well in the large binding pocket of CYPA11A1. Two strong hydrogen bond interactions were observed in the complex in which the nitrogen in the azole group forms the first hydrogen bond with Asn210 residue, whereas the oxygen atom in the same azole group forms the second hydrogen bond with Gln377 residue (Figure 2C, D). In silico mutagenesis of these two key residues, Asn210 and Gln377 to alanine followed by docking showed that Gln377 contributed more to the stability of the Posaconazole-CYP11A1 complex (Supplementary Figure S2 A-C) and resulted in elimination of hydrogen bonding interactions with both Asn210 and Gln377. Mutation of Gln377 to alanine residue has resulted in elimination of the key hydrogen bonding interaction between Gln377 and Posaconazole. The distance between azole oxygen of Posaconazole and Gln377Ala has increased from 3 Å to 6.7 Å in the docking pose after mutation and resulted in a less stable docking pose indicated by docking score (Supplementary Figure. S2 B). However, after mutating to alanine, in the absence of Asn210, Posaconazole could retain the hydrogen bonding interaction with Gln377. Posaconazole has an N-aryl piperazine backbone attached to the phenyl and azole group suitable for hydrophobic interactions. Non-polar interactions stabilizing the Posaconazole-CYP11A1 complex is contributed by amino acids such as Glu52, Tyr61, Ser59, Val57, Leu54, Phe82, Val35, Iue209, Gln356, Ile84, Ser352, Arg81, Trp87, Leu101, Thr282, Trp231, Ala286, Met201, Phe202, Gly287, Ile461, Ile351, Val353 and Phe458. Among the non-polar interactions, the terminal 2,4-difluorophenyl in the Posaconazole forms a Pi-Pi (π/π) T-shaped interaction with the Heme group, which further stabilises the complex. Val57, Ph82, Leu209, Ile84, Leu101, Ala286, Leu460 and Leu54 form Alkyl and π -Alkyl interaction between Posaconazole and CYP11A1. Stabilisation of ligand structure in the active by Van der Waals interactions was orchestrated by amino acid residues such as Glu52, Tyr61, Ser59, Val35, Ser352, Arg81, Trp87, Thr282, Trp231, Met201, Phe202, Gly287, Ile461, Ile351, Val353 and Phe458. These interactions facilitate the formation of a strong protein-ligand complex, which prevents Cholesterol from accessing the CYP11A1 active site restricting the formation of Pregnenolone, thereby forestalling the steroid biosynthesis pathway.

**Figure 2.**
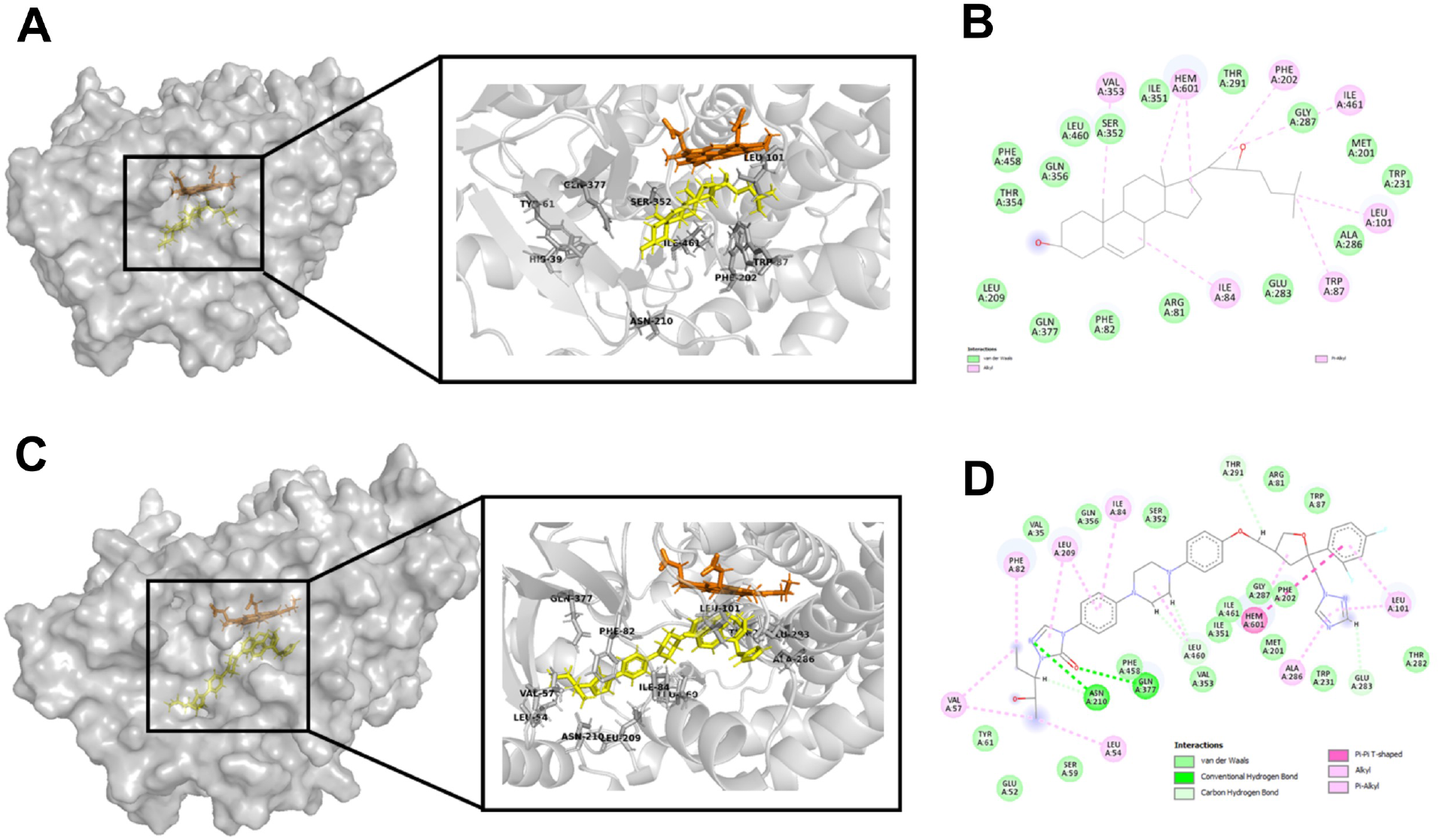
CYP11A1 crystal structure (PDB ID: 3N9Z) and docking pose of Posaconazole generated by GOLD suite. **A**. Surface structure of the CYP11A1 protein (cartoon, grey) with 22-hydroxycholesterol (stick, yellow). Inset shows the co-crystallized ligand and the key residues involved in the ligand protein interactions. Heme co-factor is shown as stick in orange. **B**. 2D interaction diagram of 22-hydroxycholestrol created using discovery studio visualizer. **C**. Surface structure of Posaconazole CYP11A1 complex. Inset shows focused image of ligand binding pocket of CYP11A1 (cartoon, grey) with Posaconazole (stick, yellow) and heme cofactor (stick, orange). The interacting residues in the protein are labelled and represented as grey sticks. **D**. 2D diagram of ligand and protein interaction created using discovery studio visualizer. Residues forming various interactions with Posaconazole (hydrogen bonds, carbon hydrogen bonds, van der Waals interaction, pi-alkyl interaction, Pi-Pi interactions and alkyl interactions) are denoted in the diagram.

### Oral administration of Posaconazole restricts experimental lung metastasis by enhancing anti-tumour immunity

Previously, it has been shown that genetic deletion or pharmacologic inhibition of Cyp11a1 improves anti-tumour immunity and restricts experimental tumour growth (Mahata et al., 2020). Therefore, to test the effect of Posaconazole *in vivo*, we administered a globally approved formulation of Posaconazole, Noxafil 40mg/ml oral suspension, to the experimental mice of metastatic melanoma at a dose of 20mg/kg body weight, daily, for 10 days. On 12^th^ day, lungs and spleens were harvested to analyse the tumour burden and immune cell phenotype. We observed a significant reduction in metastatic colonies in the Posaconazole-treated mice compared to the vehicle-treated mice (**Figure 3A**). Immunophenotyping of the lung and spleen was performed using flow cytometry. We found that Posaconazole treatment increased the number of iNOS^+^CD11b1^+^F4/80^+^ cells and reduced the number of Arg1^+^CD11b1^+^F4/80^+^ cells in the lung (**Figure 3B**). Increased iNOS expression in the macrophage suggests the increased M1 macrophages, which are known as anti-tumoural. By contrast, decreased Arg1 expression in the macrophage population suggests the decreased M2 macrophages, which are known as immunosuppressive and pro-tumoural. The observations of changes in M1:M2 because of drug treatment was further validated by decreased CD206 expression and increased CD86 expression (**Figure 3C**). In Posaconazole-treated mice, CD62L^-^CD44^+^ activated T cells were increased in frequency (**Figure 3D**). Both CD4^+^ and CD8^+^ T cells express more IFNγ and TNFα in drug treated mice compared to vehicle-treated mice (**Figure 3E, F**). We observed no significant change in the systemic level of steroids as revealed by steroid profiling of control and Posaconazole treated mice serum (**Supplementary Figure S3**).

**Figure 3.**
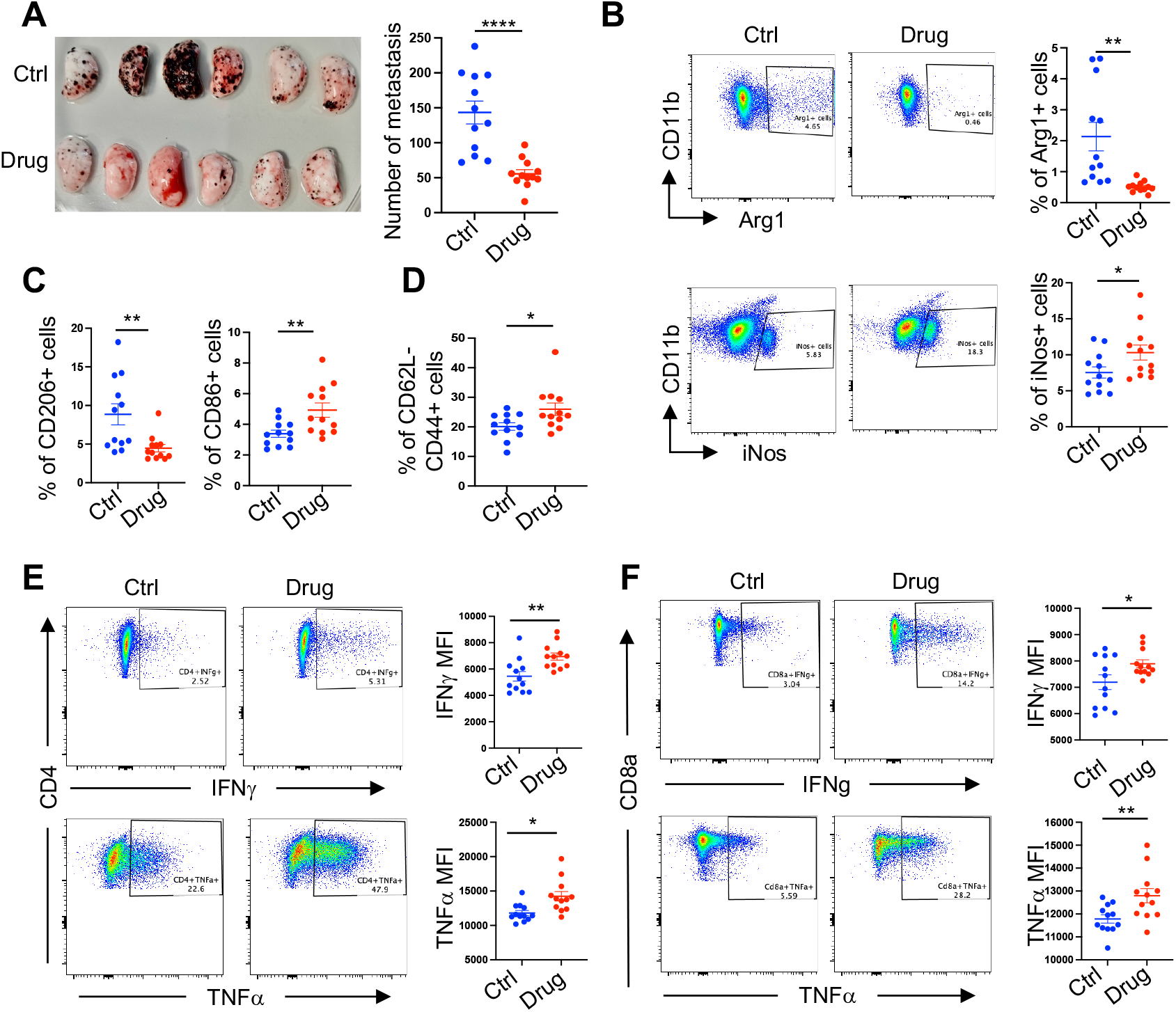
Effect of oral administration of Posaconazole in lung colonization of B16F10 melanoma cells. B16F10 cells were injected in C57BL/6 mice in the tail vein. A globally approved formulation of Posaconazole, Noxafil 40mg/ml Oral suspension, was administered to the experimental mice of metastatic melanoma by oral gavage at a dose of 20mg/kg body weight, daily, for 10 days. Lungs were harvested after 12 days. **A**. Left panel: Representative photograph of pulmonary metastatic foci produced 12 days after tail vein injection of B16-F10 cells in Posaconazole treated and control (drug and ctrl, respectively) mice. Left lobes of the lungs are shown. N = 11-12. Right panel: Graphical presentation of the numbers of lung metastatic foci. Each point represents an individual animal, error bars represent mean with s.d. Unpaired two-tailed t-test. **B-D**. Comparing immunophenotype in Posaconazole (Drug) treated and vehicle-treated (Ctrl) mice with B16-F10 cells injected intravenously. N=11-12. **B**. Representative FACS profile and respective graphical presentation of iNOS and Arg1 expression in lung-infiltrating live, singlet, CD45^+^Lin^-^ CD11b^+^ F4/80^+^ macrophages. **C**. Respective graphical presentation of CD206 and CD86 expression in lung-infiltrating live, singlet, CD45^+^Lin^-^ CD11b^+^ F4/80^+^ macrophages. **D**. Representative graphical presentation of CD62L^-^ CD44^+^ CD8 T cells. **E-F**. Representative FACS profile and respective graphical presentation of IFNγ and TNFα expression in splenic CD4 (E) and CD8 T (F) cells. *FACS Gating*: All cells > singlets > live cells > CD45^+^ > B. Lin^-^CD11b^+^F4/80^+^ > iNOS and Arg1; C. Lin^-^ CD11b^+^ F4/80^+^ D. Lin^-^ TCRβ_+_ > CD4^+^ or CD8^+^ > IFNγ or TNFα. In this figure, all error bars represent mean with s.d. P-value was calculated by unpaired two-tailed t-test. * P ≤ 0.05, ** P ≤ 0.01, *** P ≤ 0.001. N represents biologically independent animals.

These results collectively demonstrate that oral administration of Posaconazole effectively restricts experimental lung metastasis by reinvigorating anti-tumour immunity in vivo. The observed changes in immune cell composition and function suggest a shift towards a more favourable, tumour-suppressive microenvironment. This study provides compelling evidence for the potential of Posaconazole as an immunomodulatory agent in cancer therapy, warranting further investigation into its clinical applications.

## Discussion

Immunosuppression and cancer immune evasion is a therapeutic challenge in developing effective immunotherapy (Hegde and Chen, 2020). Based on mice model-based experimental evidence, it has been proposed that targeting CYP11A1 may reinstate anti-tumour immunity (Acharya et al., 2020; Mahata et al., 2020). Therefore, in this study, we aimed to identify a new inhibitor of CYP11A1 and test its anti-tumour efficacy.

The study successfully employed a drug repurposing approach to identify Posaconazole as a potent inhibitor of CYP11A1, outperforming the established inhibitor Aminoglutethimide. The in silico structure based virtual screening of 7,190 compounds from the ZINC15 database, followed by rigorous in vitro validation, led to this discovery. This approach not only accelerates the drug discovery process but also leverages existing safety and pharmacokinetic data, potentially expediting clinical translation.

The superior inhibitory effect of Posaconazole can be attributed to its molecular structure and interactions within the CYP11A1 binding pocket. Posaconazole’s larger size (700.8 g/mol) compared to Aminoglutethimide (232.28 g/mol) allows for better occupancy of the CYP11A1 binding cavity. Unlike Posaconazole, Aminoglutethimide does not display any hydrogen bonding interactions with any of the key residues present in the active site. The hydrophobic ring structures of Posaconazole align well with the hydrophobic regions of the active site, mimicking the natural ligand cholesterol. In the docked conformation of CYP11A1-Posaconazole complex, the terminal phenyl-2-hydroxypentan-triazolone group is positioned near to the surface away from the heme group. The presence of this group is critical for the stability of the complex by forming polar interactions in the active site. Posaconazole forms two critical hydrogen bonds: one between its nitrogen atom and Asn210, and another between its oxygen atom and Gln377, present in the triazolone group. In the docked conformation of CYP11A1-Posaconazole complex, the terminal phenyl-2-hydroxypentan-triazolone group is positioned near to the surface away from the heme group. The presence of this group is critical for the stability of the complex by forming polar interactions in the active site. Leu101 forms alkyl interactions with the 1,2,4 triazole group and p-alkyl interactions with the 2,4-difluorophenyl ring. The 2,4-difluorophenyl ring of Posaconazole engages in pi-pi interactions with the heme cofactor. Another important residue involved in positioning native ligand cholesterol’s C22 atom near to the heme cofactor is Ser352, along with the hydrogen-bonded water molecule (Strushkevich et al., 2011). The Ser352 residue forms van der Waals interaction to stabilise the protein-ligand complex in the Posaconazole docked conformation. These interactions collectively stabilize the Posaconazole-CYP11A1 complex, effectively blocking cholesterol access to the active site and inhibiting pregnenolone synthesis.

The dose-dependent inhibition of pregnenolone synthesis by Posaconazole suggests its potential application in cancer immunotherapy. Our in vivo studies demonstrated that oral administration of Posaconazole restricts tumour growth in a mouse model of metastatic melanoma by reinvigorating anti-tumour immunity. Posaconazole treatment increased anti-tumoural M1 macrophages and decreased pro-tumoural M2 macrophages in the lung. The drug increased the frequency of activated T cells and enhanced IFNγ and TNFα expression. These findings suggest that Posaconazole can reshape the tumour microenvironment to favour anti-tumour immune responses.

Posaconazole provides an immediate benefit as a research tool to study steroidogenesis, offering a means to inhibit the first and rate-limiting step of the steroid biosynthesis pathway. Beyond cancer, Posaconazole’s CYP11A1 inhibition could be explored in other conditions characterized by elevated steroidogenesis, such as peanut allergy. As an FDA-approved antifungal medication, Posaconazole’s safety profile and pharmacokinetics are well-established, potentially accelerating its translation to new therapeutic applications. The study provides valuable insights into the structural basis of CYP11A1 inhibition, which could guide future drug design efforts targeting this enzyme.

### Limitations of the study

While our study demonstrates the potential of Posaconazole as a CYP11A1 inhibitor, several limitations present opportunities for further research. Expanding the compound library beyond approved drugs could potentially identify even more potent CYP11A1 inhibitors. Structural biology investigations, such as co-crystallisation of Posaconazole with CYP11A1, along with introduction of point mutations in the key CYP11A1-Posaconazole interacting sites would provide valuable insights into their precise interactions. In vitro studies involving mutations of the predicted interacting residues could further validate our in silico observations. The molecular mechanisms underlying Posaconazole’s immunomodulatory effects require further elucidation. Potential off-target effects of Posaconazole were not extensively explored in this study. An in vivo comparative study with aminoglutethimide, approved immunotherapies, such as checkpoint inhibitors, or other cancer drugs would be an important future direction. These limitations provide opportunities for future research to fully understand Posaconazole’s potential as a cancer immunotherapy agent, underscoring the promising foundation laid by our study for exploring its role in immunomodulation and cancer prevention through targeting immune cell-mediated steroidogenesis.

In conclusion, this study identifies Posaconazole as a promising CYP11A1 inhibitor with potential applications in cancer immunotherapy and other conditions characterized by elevated steroidogenesis. The immediate benefit lies in its use as a research tool to study steroidogenesis, while its clinical potential warrants further preclinical and clinical investigation.

## Materials and Methods

### Mice

In this investigation, all mice were handled and cared for following the stringent guidelines set forth by the UK Animals in Science Regulation Unit’s Code of Practice for the Housing and Care of Animals Bred, Supplied, or Used for Scientific Purposes, and the Animals (Scientific Procedures) Act 1986 Amendment Regulations 2012. All experimental protocols were conducted under the authorization of a UK Home Office Project License (PPL P0AB4361E) and received the necessary approval from the local institute’s Animal Welfare and Ethical Review Body, ensuring compliance with ethical standards and animal welfare considerations.

The sample size was determined according to our previous experience and a priori power analysis (G^*^Power). Housing condition of Gurdon animal facility: all mice used in this study were maintained in specific pathogen free unit on 12 hours light and 12 hours dark cycle. The mice were genotyped by Transnetyx. In this study we used all male mice aged between 8 to 12 weeks.

### B16-F10 melanoma metastasis model and Posaconazole treatment

B16-F10 melanoma cell line was purchased from American Type Culture Collection (ATCC) and cultured in Dulbecco’s Modified Eagle medium (DMEM, Life Technologies), supplemented with 100 U/mL penicillin/streptomycin and 10% foetal bovine serum (FBS; Life Technologies, Invitrogen). For metastasis, 5 × 10^5^ B16-F10 cells in a volume of 0.1 ml PBS were injected intravenously into the tail vein of wild-type (WT) C57BL/6 mice. Mice received 200μL Posaconazole (Noxafil, 40mg/ml oral suspension) by oral gavage at 20 mg/kg body weight, diluted in water, once daily (Francisco et al., 2015; Salas et al., 2012; Ullmann et al., 2007). The first dose was administered concurrently with cell injection and continued for 10 days. The vehicles were administered orally by gavage in control mice. After 12 days, animals were killed and perfused the lung with 10 ml PBS, and tissues were collected for analysis. The number of B16-F10 colonies on all five lobes of the lung was counted macroscopically.

### Steroid profiling of mice serum

Multiple steroids were measured in serum samples following the method described in (Lahti-Pulkkinen et al., 2025) by targeted liquid chromatography tandem mass spectrometry (LC-MS/MS). Briefly, serum was enriched with a mixture of isotopically labelled steroids as internal standards and steroids were extracted by supported liquid extraction on an Extrahera liquid handling robot by Biotage (Uppsala, Sweden). The extract was subject to chromatographic separation on an Acquity I-Class UPLC by Waters (Wilmslow, UK) fitted with a Kinetex C18 (150 × 2.1 mm; 2.6 um) chromatography column by Phenomenex (Macclesfield, UK). Steroids were separated using a 16-minute gradient mobile phase consisting of water and methanol, both with 0.05 mM ammonium fluoride, with a flow rate of 0.3 mL/min and a column temperature of 50C. Separated steroids were analysed on a QTrap 6500+ by AB Sciex (UK) using multiple reaction monitoring in both positive and negative ion mode. Concentrations of steroid in the sample were calculated using MultiQuant software by AB Sciex (Macclesfield, UK) as described in (Denham et al., 2024).

### ‘Lung tissue processing for metastatic melanoma analysis

The lungs from metastatic melanoma bearing mice were mechanically dissociated and were then subjected to enzymatic digestion using a mixture of 1 mg/ml collagenase D (Roche), 1 mg/ml collagenase A (Roche), and 0.4 mg/ml DNase I (Sigma) in IMDM media containing 10% FBS, all incubated at 37 °C for 30-40 minutes. To halt collagenase activity, EDTA (5mM) was introduced to all samples. The digested tissues were finally strained through 70μm cell strainers (Falcon) to ensure uniformity in sample preparation.

### Flow Cytometry

Cells prepared for flow cytometric analysis were subjected to a 4-hour incubation with PMA (50 ng/ml) and Ionomycin (500 ng/ml), followed by the addition of Monensin (Biolegend) to the culture for the last 3 hours. The established protocols for surface staining and intracellular cytoplasmic protein staining by eBioscience were diligently followed. LIVE/DEAD Fixable Dead Cell Stain (Thermo Fisher) was employed to stain the single-cell suspensions, which were subsequently fixed using the eBioscience IC fixation buffer. For intracellular cytokine staining, the cells were fixed and permeabilized using eBioscience IC Fixation buffer and 1x permeabilization buffer, respectively. Fluorescent dye-conjugated antibodies were then applied to stain the cells. After a thorough wash with 3% PBS-FCS, the cells were ready for analysis using the Cytek Aurora (5L) flow cytometer. FlowJo v10.2 software facilitated the data analysis.

### T helper cell culture

Splenic CD4^+^ T helper cells from C57BL/6 mice were purified using the CD4^+^ T Cell Isolation kit (Miltenyi Biotec). 100,000 CD4^+^ T cells were seeded in 200μL on round bottom 96 well tissue culture plate anti-CD3e (2 μg/ml, clone 145-2C11, eBioscience) and anti-CD28 (3 μg/ml, clone 37.51, eBioscience) coated plates in Iscove’s Modified Dulbecco’s Medium containing 10% charcoal-stripped serum (without Phenol red), β-mercaptoethanol (55 μM) and recombinant murine IL2 (10 ng/ml, R&D Systems). Charcoal-stripped serum was used to reduce the background of lipid and steroids in the Pregnenolone ELISA. To polarise the cells toward T helper 2 (Th2), recombinant murine IL4 (10 ng/ml, R&D Systems), neutralising anti-IFNγ antibody (10 μg/ml, clone XMG1.2, eBioscience) and anti-IL12 antibody (10 μg/ml) were added. The Th2 cells were transferred from the activation plates to resting plates after 72 hours (in a 5% CO2 incubator). The drugs were then added to the Th2 cells with fresh media (serially diluted), and the control wells received DMSO or vehicle. After 48 hours, the plates were centrifuged at 400 X g for 5 min to collect cell supernatant for Pregnenolone quantification using ELISA. The cells in each well were used to assess the effect of drug treatment on cell viability.

### H295R cell culture and Pregnenolone ELISA

H295R cells were cultured in a 1:1 mixture of DMEM/Ham’s F-12 nutrient medium with phenol red, supplemented with 2.5% Nu-serum, 1% ITS+ Premix, and 1% penicillin-streptomycin at 37°C in a humidified atmosphere (95% air, 5% CO2).

### Y1 cell culture

Y1 cells were cultured in Ham’s F-12K medium, supplemented with 2.5% foetal bovine serum, 15% horse serum, and 1% penicillin-streptomycin.

### Cytotoxicity assay

For the analysis cytotoxicity of the drugs, Th2 cells treated with compounds for 48 hours were stained with a solution composed of 80 µg/ml Acridine Orange (AO) and 40 µg/ml DAPI (Chemometec). The percentage of viable and dead cells were counted using NucleoCounter NC-250 automated cell counter (Chemometec). Percentage viability of cells treated with drugs with respect to the viability of control cells were calculated for identification of cytotoxicity of candidate drugs under study.

### Quantitative Pregnenolone ELISA

*In vitro* differentiated Th2 cells from the 4^th^ day were cultured in the presence of our chosen drugs for 48 hours at a density of 1 × 10^6^ cells per mL in 200μL volume in absence of any further activation. After this period, the plates were spun for 5 minutes at 400 X g at room temperature, and the supernatant was collected for ELISA analysis. H295R and Y1 cells between passages 5 and 10 were seeded in 6-well plates (2 mL medium per well) and allowed to adhere for 24 hours. The cells were then treated with either vehicle (DMSO) or Posaconazole at concentrations ranging from 0.78 μM to 25 μM in the IMDM medium without phenol red and supplemented with 10% charcoal-stripped FBS and 1% penicillin-streptomycin. After 48 hours of incubation at 37°C in a humidified atmosphere (95% air, 5% CO2), the culture supernatant was collected for pregnenolone ELISA analysis. Pregnenolone concentration of the collected samples was quantified using the pregnenolone ELISA kit (Abnova) following the manufacturer’s instructions. Absorbance was measured at 450 nm and analysed using GraphPad Prism and Excel.

### Protein Preparation

The available structures of the CYP11A1 were downloaded from PDBe (Armstrong et al., 2020). Water molecules were deleted and the hydrogen atoms were added before docking using CCDC GOLD suite(Jones et al., 1997). The ligand 22-hydroxy cholesterol and the iron-sulfur clusters present in the crystal structure removed before docking. The Heme cofactor within the active site of the CYP11A1 contributes to the ligand bindings, and therefore it was retained during protein preparation for virtual screening. *In silico* mutagenesis of the protein was done using PyMol.

### Ligand preparation

Three libraries of compounds were downloaded from the ZINC15 database (Sterling and Irwin, 2015) (https://zinc15.docking.org), namely the FDA subset, World subset and Investigational only subset. A KNIME workflow (**Supplementary Figure S4**) using RdKit nodes (RDKit: Open-source cheminformatics (https://www.rdkit.org) were used to generate 3D conformers of the ligands. Missing hydrogen atoms were added prior to the conversion. Finally, CCDC conformer generator was used to make a single conformer of each ligand.

### Virtual Screening

A brief overview of the screening methodology is represented in **Figure 1A**. CCDC GOLD suite was used for the virtual screening. The prepared structure of the target protein was used to perform a preliminary docking experiment using GOLD, where the crystallised ligand or Cholesterol was redocked with the protein. Consensus scoring was used to determine the best scoring function to be used for further docking runs. CHEMPLP provided the best performance. This docking configuration was used to perform virtual screening with all three libraries of the compounds.

Selection of compounds for experimental validation: The docking results of all three subset libraries were combined, and the top 100 poses with the highest docking scores were shortlisted for further analysis. DataWarrior(Sander et al., 2015) was used to cluster the 100 compounds with similarity cut-off criteria 0.8. The top-scored one was taken for further analysis in the clusters with more than one compound. A total of 69 compounds were taken for visual analysis after clustering. PyMol and Discovery studio visualizer were used for visual analysis and further evaluation of the docking poses. CCDC Mogul(Bruno et al., 2004; Cottrell et al., 2012) was used to assess ligand geometries.

### Statistical Analysis

The statistical analysis was performed using GraphPad Prism Software. The data were analysed by either one-way or two-way ANOVA. The Dunnet multiple comparison test was employed for one-way ANOVA, and for two-way ANOVA, Sidak multiple comparison test was used.

Unpaired two-tailed t-test has also been applied in some experiments. The p values < 0.05 were considered significant. The values were presented as mean ± standard deviation.

## Author contributions

**Conceptualization:** BM, AM. **Methodology:** BM, SKS, JP. **Investigation:** JP (wet lab, in vivo, in vitro), SKS (in silico, in vitro), MZ, BB (preliminary work), BZ, YK (in vivo), NZMH (Steroid analysis). **Data Curation & Visualization:** JP, SKS, NZMH (Steroid detection). **Writing – Original Draft:** SKS, JP, BM. **Writing – Review & Editing:** All authors. **Supervision:** BM (overall), RR (immunology), KO (signalling), NZMH (Steroid detection), AM (computational). **Funding Acquisition:** BM. **Project Administration:** BM

## Acknowledgement

This study is supported by CRUK Career Development Fellowship (RCCFEL\100095), NSF-BIO/UKRI-BBSRC project grant (BB/V006126/1) and MRC project grant (MR/V028995/1). JP is supported by NSF-BIO/UKRI-BBSRC project grant (BB/V006126/1) and MRC project grant (MR/V028995/1).

