## Supplementary Figures for "Drug repurposing reveals Posaconazole as a CYP11A1 inhibitor enhancing anti-tumour immunity"

**Supplementary Figure S1**

**A**

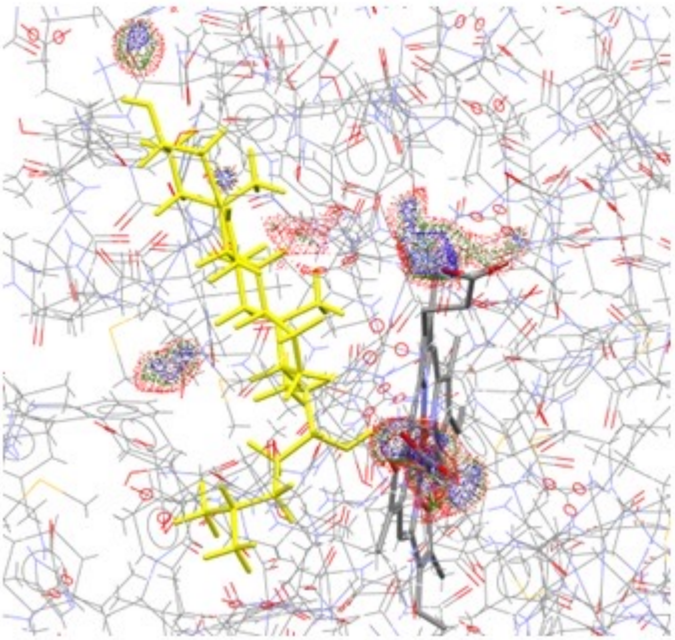

**B**

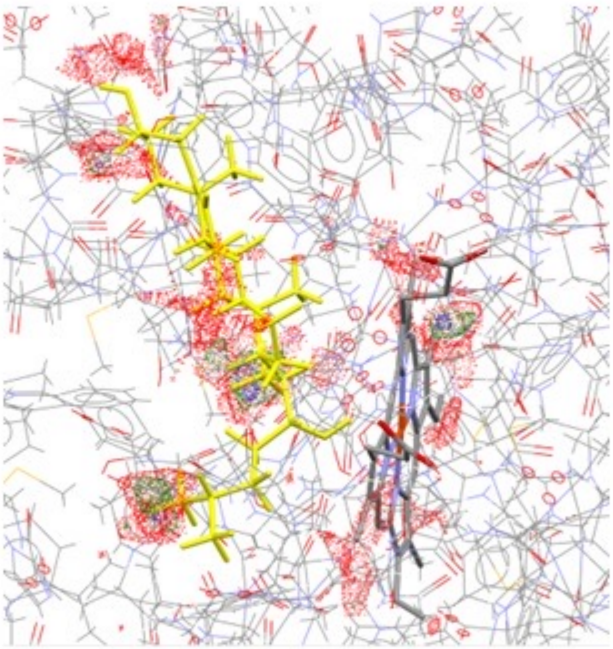

**C**

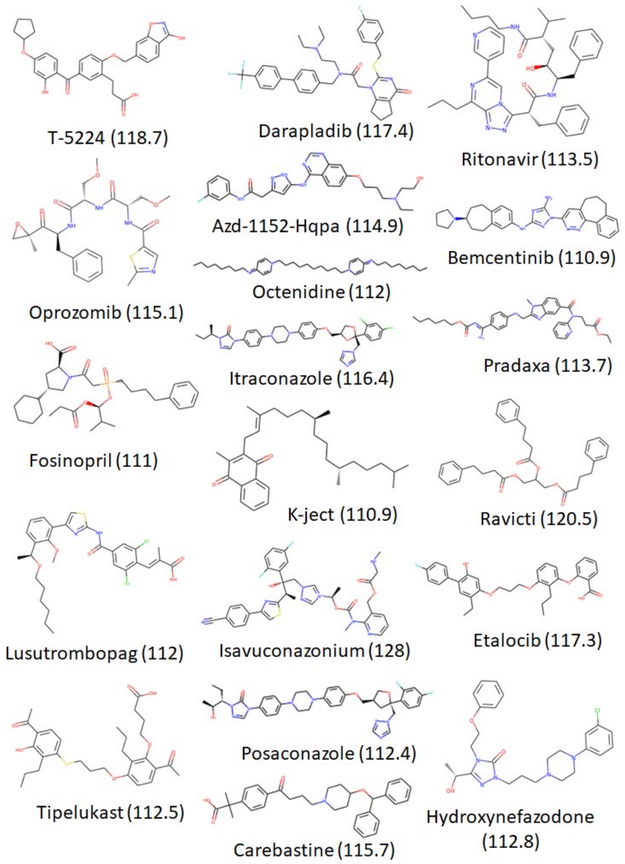


**D**

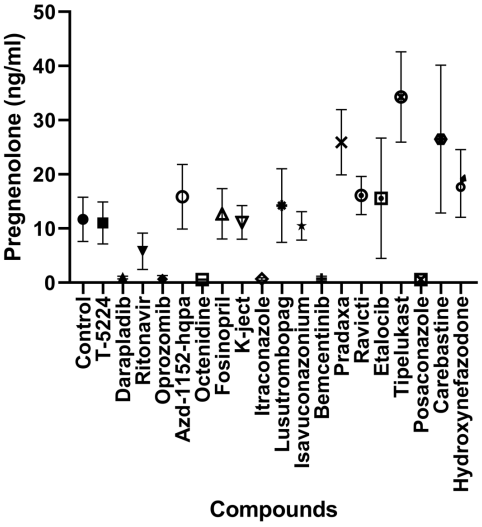


**E**

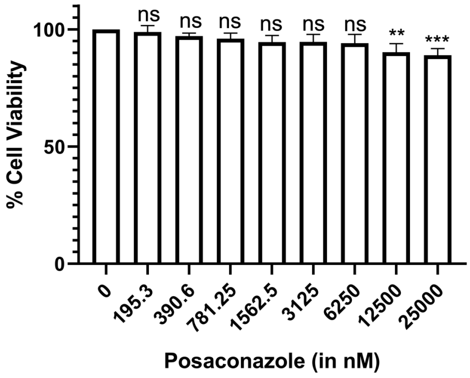

**Supplementary Figure S2**

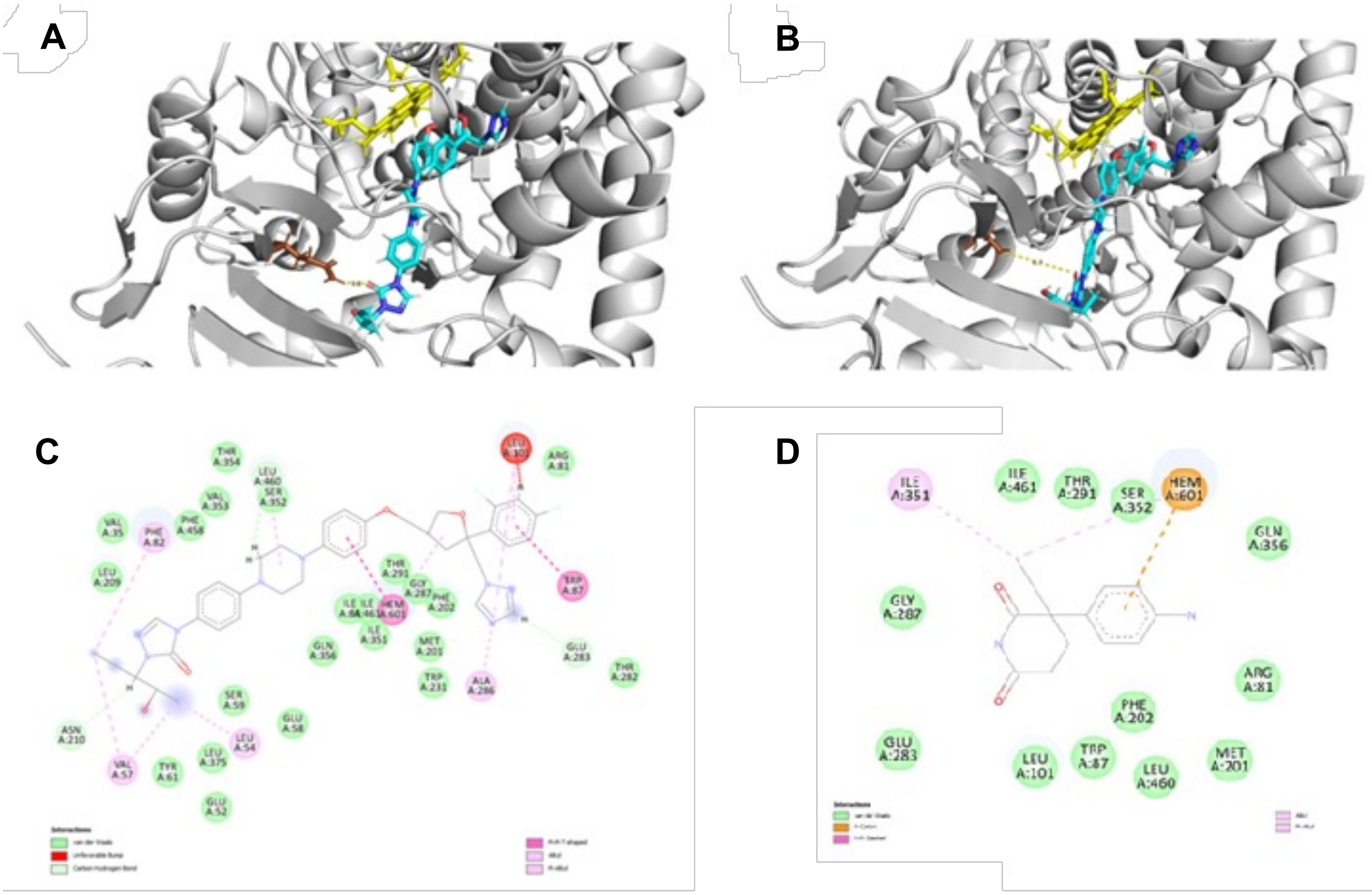

**Supplementary Figure S3**

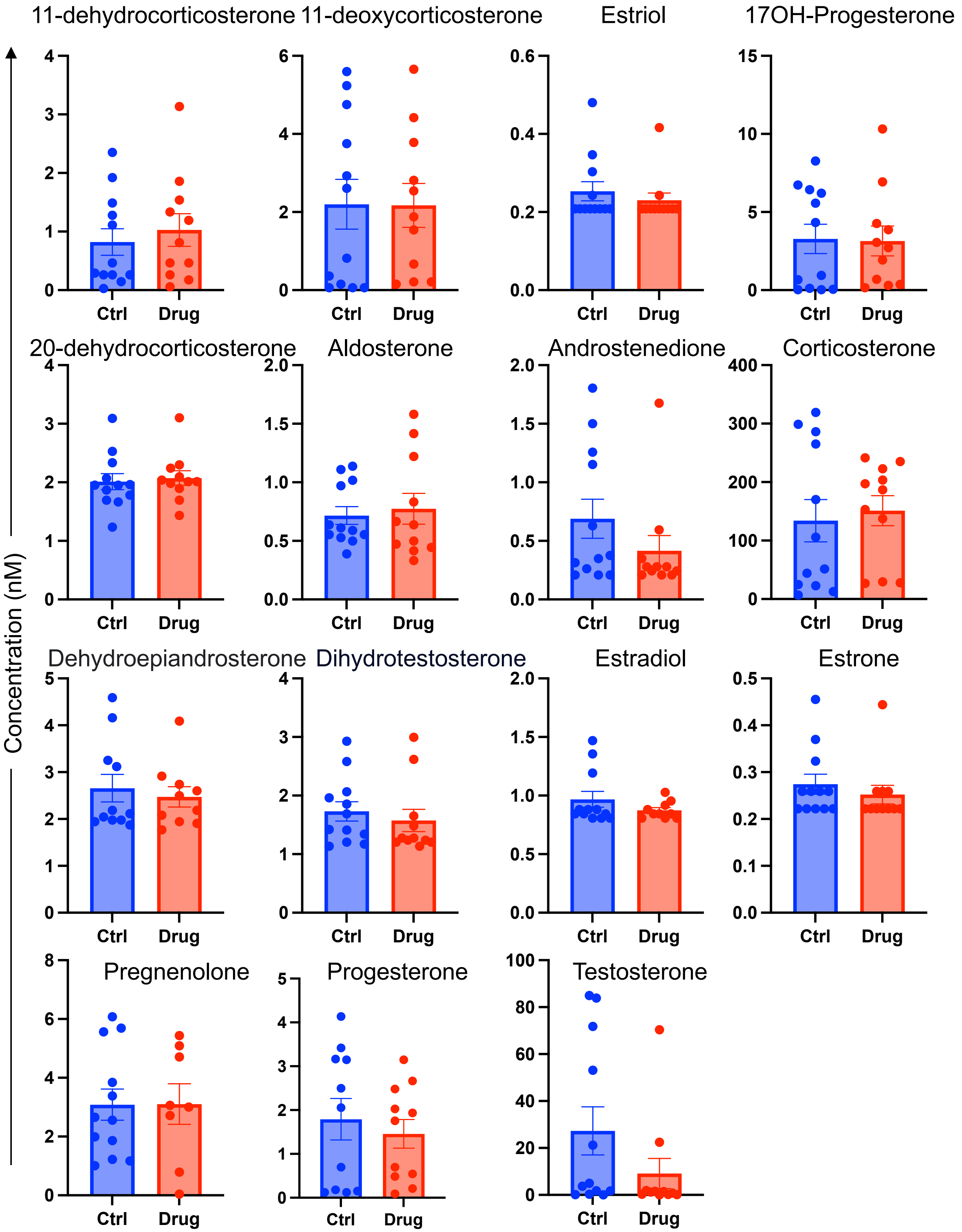

**Supplementary Figure S4**

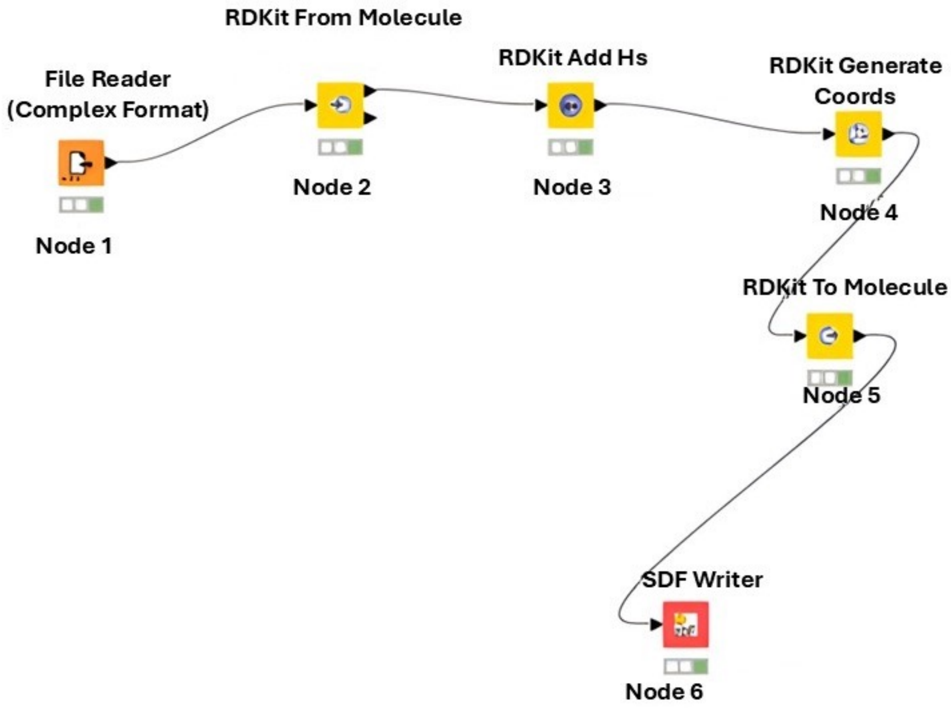
